# bulk2scDiff: A Pseudobulk-Conditioned Diffusion Model for Bulk-to-Single-Cell RNA-Seq Generation

**DOI:** 10.64898/2026.08.20.745960

**Authors:** Jiedan Xiao, Andreas Raue

## Abstract

Bulk RNA sequencing remains the predominant profiling strategy for large clinical cohorts, but it aggregates transcriptional signals across cell populations, thereby masking the underlying cellular heterogeneity. Inferring this heterogeneity from existing bulk transcriptomic data could extend large cohort-based studies that have already been profiled, but constitutes an underdetermined inverse problem, as one bulk profile can be compatible with multiple underlying cellular populations. Existing computational deconvolution methods address this problem primarily by estimating cell-type proportions or cell-type-averaged expression profiles rather than resolving expression at the level of individual cells. Here, we present bulk2scDiff, a proof-of-concept conditional diffusion framework that reformulates bulk-to-single-cell inference as conditional generation of single-cell expression profiles from pseudobulk transcriptomic input. We evaluated bulk2scDiff on two cancer single-cell RNA sequencing datasets, breast cancer and acute myeloid leukemia, where pseudobulk profiles were derived from the single-cell data and used as conditioning inputs, with the matched single-cell populations providing ground truth for controlled evaluation. Across both cases, bulk2scDiff closely reconstructed populations from training samples and generated biologically coherent single-cell populations for held-out samples, generalizing most consistently to recurrent immune features. A pseudobulk-swap control further confirmed sample-specific conditioning, with each sample’s corresponding pseudobulk yielding the closest agreement with its observed population in nearly all cases. Overall, our work establishes the feasibility of conditional diffusion for generating single-cell populations from pseudobulk transcriptomic profiles, providing a foundation for future evaluation with clinical bulk RNA sequencing data.

## Introduction

Single-cell transcriptomics has transformed the study of complex tissues, tumor ecosystems, and cellular heterogeneity by resolving gene expression at single-cell resolution (1,2). In oncology and immunology, single-cell RNA sequencing (scRNA-seq) has enabled the characterization of diverse cellular states, rare subpopulations, lineage trajectories, and interactions between malignant and non-malignant compartments (3,4). These capabilities have advanced our understanding of disease progression, therapeutic response, and resistance mechanisms. However, large clinical cohorts are still profiled predominantly with bulk transcriptomic assays, which are cheaper, experimentally simpler, compatible with archived material (5), and extensively available in biobanks, such as The Cancer Genome Atlas (TCGA) (6) and the Genotype-Tissue Expression (GTEx) Program (7). Bulk measurements aggregate transcriptional signals across heterogeneous cellular populations and therefore obscure the cellular heterogeneity underlying the observed sample-level profile (8).

Recovering cellular populations from such aggregate transcriptomic measurements constitutes an inherently underdetermined inverse problem: multiple distinct cellular populations can be compatible with the same or similar bulk expression profile. Existing computational approaches address this problem through deconvolution, which uses reference expression profiles to deconstruct the mixed transcriptional signal in bulk data. Methods such as CIBERSORTx (9) and MuSiC (10) model bulk expression as a mixture of cell-type-specific signatures, allowing the relative abundance of different cell types to be estimated. Beyond cell-type proportions, CIBERSORTx (9) can additionally infer cell-type-averaged expression profiles from bulk transcriptomic data. More recent methods have incorporated deep learning into transcriptomic deconvolution. TAPE (11) uses an autoencoder trained on paired bulk and single-cell reference data to estimate cellular composition, whereas DECODE (12) formulates deconvolution as a deep-unfolded matrix factorization initialized from a reference signature matrix. Despite these methodological advances, deconvolution approaches generally reduce the underlying cellular population to summary quantities, rather than generating individual cellular transcriptomes that represent the heterogeneity within a sample. Their dependence on reference-defined cellular information can also limit the representation of context-specific states that are poorly captured by the available reference (8).

The underdetermined nature of transcriptomic aggregation motivates a different formulation of bulk-to-single-cell inference. Rather than treating an aggregate profile as uniquely specifying one underlying cellular population, the problem can be formulated probabilistically, with cellular populations generated conditional on the observed sample-level transcriptomic profile. Generative modeling provides a natural framework for this formulation because generative models learn distributions from which individual observations can be sampled rather than returning only a deterministic point estimate. Previous approaches have generated single-cell transcriptomic data using variational autoencoders (13), generative adversarial networks (14), and flow-matching frameworks (15). Diffusion models represent another class of generative approaches, originally developed for image synthesis by learning data distributions through the reverse of a gradual noising process (16), and subsequently adapted to single-cell transcriptomics, such as scDiffusion (17). Existing single-cell generative models, however, have primarily focused on generation conditioned on intrinsic annotations or experimental variables, such as cell type, perturbation, or developmental stage. The use of continuous, sample-level transcriptomic profiles as conditioning information for generating corresponding single-cell populations remains comparatively unexplored.

Here, we present bulk2scDiff, a conditional diffusion framework that reformulates bulk-to-single-cell inference as conditional generation of single-cell expression profiles from aggregate transcriptomic input. As a proof of concept, we use pseudobulk profiles derived from scRNA-seq as conditioning inputs, with the matched single-cell populations providing ground truth for controlled evaluation. This design allows us to test whether sample-level aggregate expression contains sufficient information to condition the generation of biologically coherent cellular populations before extending the framework to independently measured bulk RNA-seq. We evaluate bulk2scDiff on two independent cancer datasets of breast cancer (BRCA) and acute myeloid leukemia (AML) (18,19), and assess both reconstruction of populations from training samples and generalization to held-out samples. In this formulation, the conditioning profile constrains generation without being assumed to uniquely determine a single underlying cellular configuration.

By framing the relationship between aggregate and single-cell transcriptomes as conditional generation, bulk2scDiff provides a complementary perspective to conventional deconvolution. Rather than replacing established approaches for estimating cellular composition, the framework investigates whether aggregate transcriptomic measurements can condition the generation of cellular-resolution expression distributions. The present pseudobulk-based evaluation establishes a controlled first step toward this goal; determining whether the approach can recover meaningful cellular heterogeneity from independently measured clinical bulk RNA-seq will require validation on larger matched bulk and single-cell cohorts.

## Results

Bulk2scDiff represents single-cell profiles in a latent space defined by a pretrained SCimilarity encoder (20), while sample-level pseudobulk profiles are independently encoded into a separate conditioning embedding. During training, diffusion progressively adds noise to the single-cell latent space, and a denoising network learns to reverse this process, conditioned on the pseudobulk embedding via feature-wise linear modulation (FiLM) (21). During generation, the model starts from random noise and denoises under a given pseudobulk profile, after which the resulting latents are decoded by SCimilarity (20) into synthetic single-cell expression profiles (Figure 1a).

**Figure 1:**
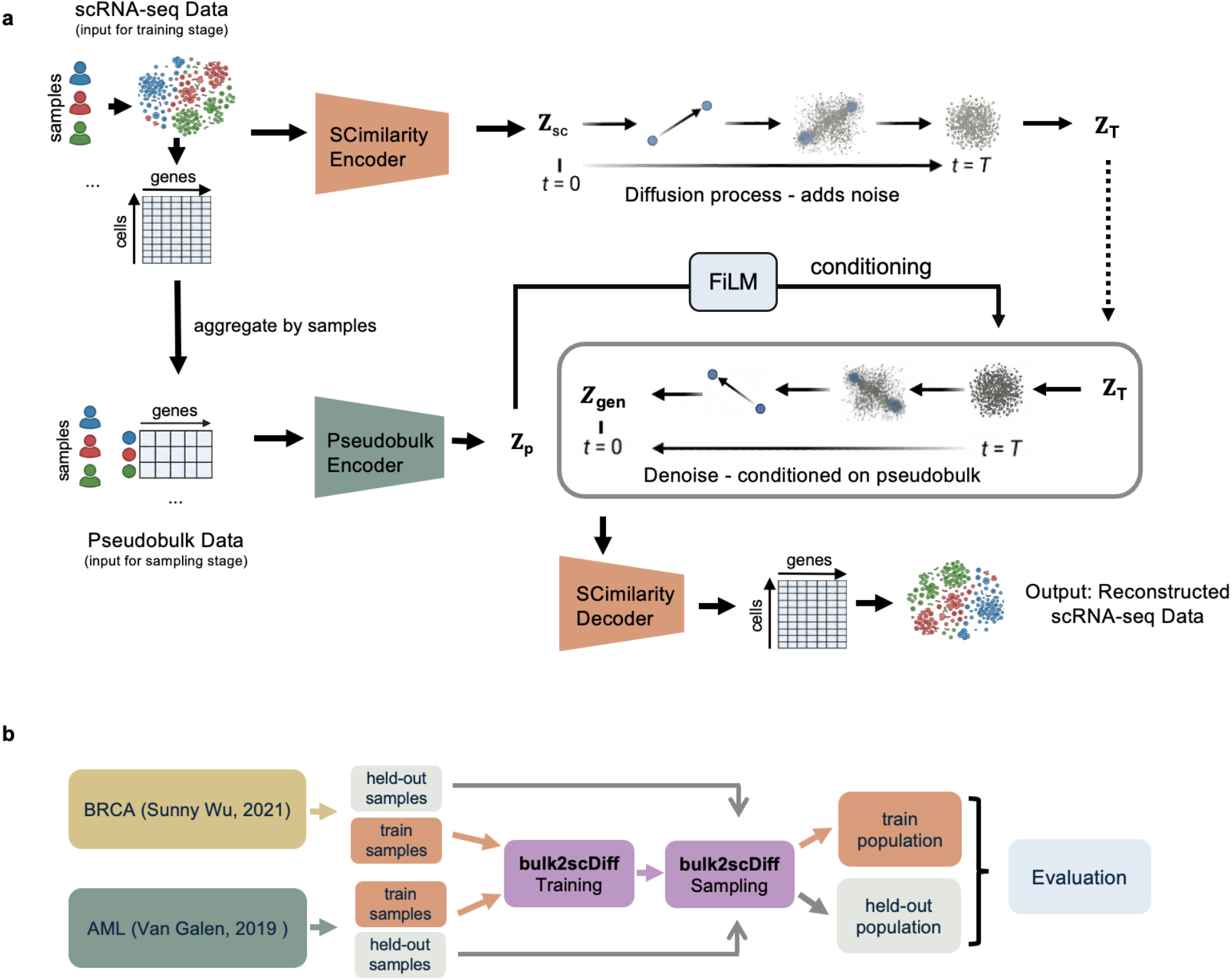
Model architecture and evaluation workflow. a) Schematic of the bulk2scDiff architecture. Single-cell profiles are encoded into a latent representation, while pseudobulk profiles are encoded separately and used to condition the latent diffusion model during denoising. b) Sample-level evaluation workflow. For each dataset, samples are split into training samples, used for model optimization, and held-out samples, reserved for independent evaluation.

We evaluated whether bulk2scDiff could learn this conditional mapping from pseudobulk transcriptomic profiles to sample-specific single-cell distributions using two cancer single-cell RNA-seq datasets, a breast cancer cohort (19) and an AML cohort (18). Samples in each dataset were split at the sample level into training and held-out sets (Table 1). Held-out samples were excluded from diffusion training and therefore represented unseen sample-level conditioning inputs during generation. The evaluation workflow is summarized in Figure 1b.

**Table 1:** Dataset overview. Training and held-out sets were defined at the sample level.

| Datasets | Samples | Training | Held-out | Cells (Training / Held-out) | Notes |
| --- | --- | --- | --- | --- | --- |
| BRCA (Wu et al., 2021) | 26 | 21 | 5 | 81,177 / 18,349 | 11 ER+, 5 HER2+, 10 TNBC samples (subtype-balanced split) |
| AML (van Galen et al., 2019) | 41 | 32 | 9 | 29,662 / 8,146 | 35 AML patient samples and 6 normal bone marrow samples; MUTZ3 and OCI-AML3 cell-line samples excluded |

### Global transcriptomic organization and cell-type structure

We first examined whether pseudobulk-conditioned generation recovered the global transcriptomic organization of the real single-cell datasets. In both the breast cancer and AML cohorts, real cells formed the expected major cell-type compartments in Uniform Manifold Approximation and Projection (UMAP) space, including malignant and non-malignant cellular populations (Figure 2a). Generated cells broadly overlapped with these populations, recapitulating the major hematopoietic and leukemic regions in AML and the major immune and tumor-associated regions in breast cancer (Figure 2b). Most generated cells mapped to regions occupied by real cells, whereas a small number of real-cell clusters were not fully recovered. These underrepresented clusters primarily represented sample-specific malignant populations rather than shared immune populations, suggesting that recurrent cellular programs were more consistently reproduced than highly sample-specific malignant states. Thus, despite the distinct cellular composition of the two cohorts, bulk2scDiff recovered the major transcriptomic organization observed in both hematologic and solid tumor datasets, while showing reduced coverage of sample-specific malignant populations.

**Figure 2:**
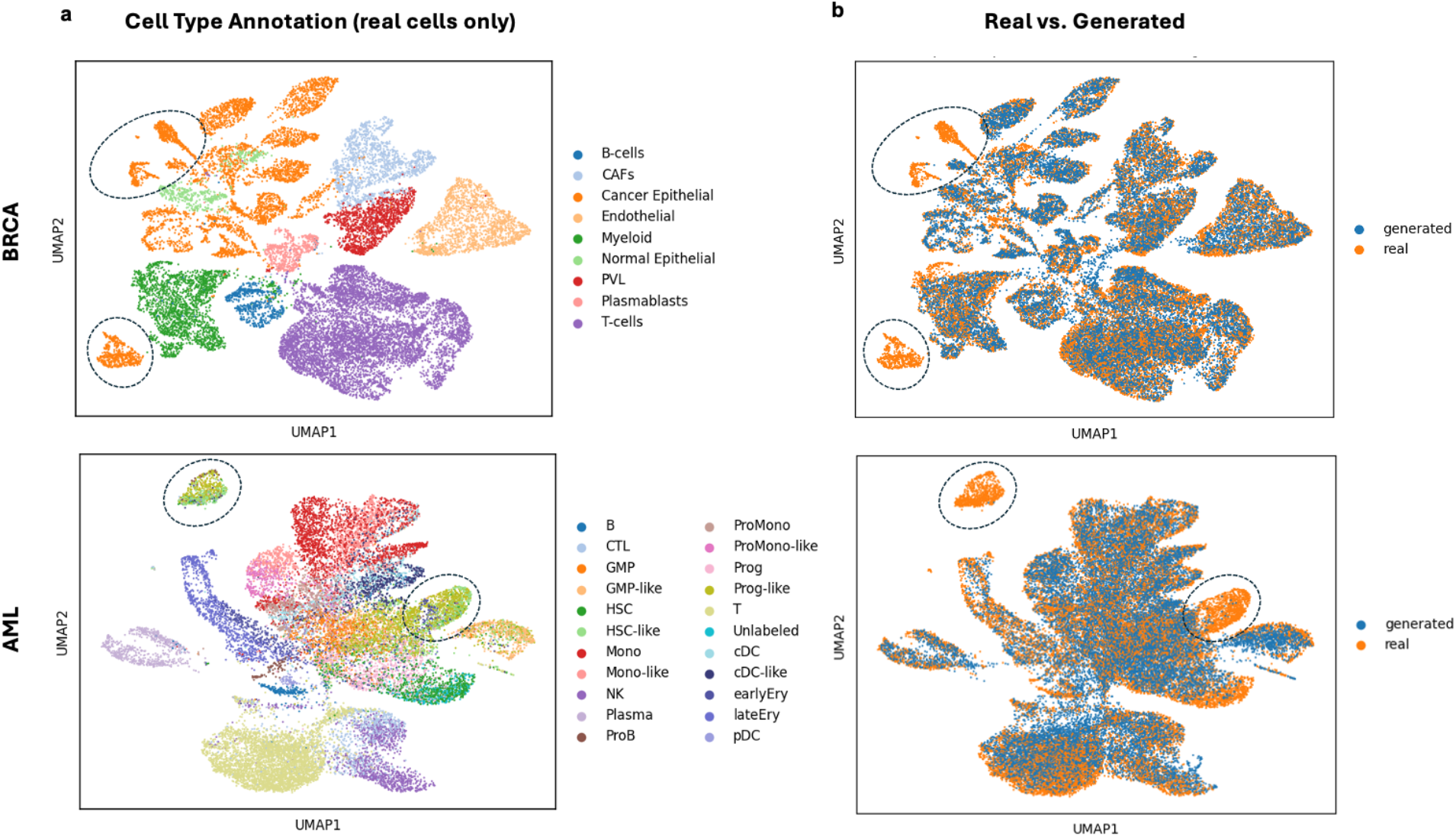
Global performance and cell type annotation. For the BRCA and AML datasets, UMAP embeddings show the structure of the real single-cell RNA-seq data colored by cell type (a) and the overlap between real and generated cells in the same embedding space (b).

To assess how this global agreement extended beyond the samples used for model training, we next examined the global embedding separately for training and held-out samples (Supplementary Figure 1). Generated cells from training samples showed strong overlap with their corresponding real-cell distributions, whereas those from held-out samples still covered most major real-cell populations but showed less complete overlap, particularly for sample-specific malignant cell regions. This contrast shows that bulk2scDiff robustly reconstructs the cellular distributions under observed sample-level conditions, while generalization to unseen samples is more limited for sample-specific malignant populations.

### Reconstruction of immune marker gene expression

We next asked whether the generated cells also preserved biologically meaningful gene-expression patterns, evaluating immune and malignant marker genes separately given their distinct patterns of reconstruction under pseudobulk conditioning. We first examined immune marker gene expression in the breast cancer dataset.

For immune populations, generated cells reproduced the spatial distribution and expression strength of canonical marker genes across the UMAP embedding. *CD3D* expression localized to T-cell regions, *CD68* to macrophage regions, and *MS4A1* to B-cell regions in the real data, with corresponding patterns closely mirrored in generated cells from both training and held-out samples (Figure 3). The agreement extended to housekeeping genes: *ACTB*, representing a broadly expressed gene, and *HPRT1*, representing a lower-expression housekeeping gene, exhibited similar expression intensity and spatial distribution in both training and held-out samples (Supplementary Figure 2).

**Figure 3:**
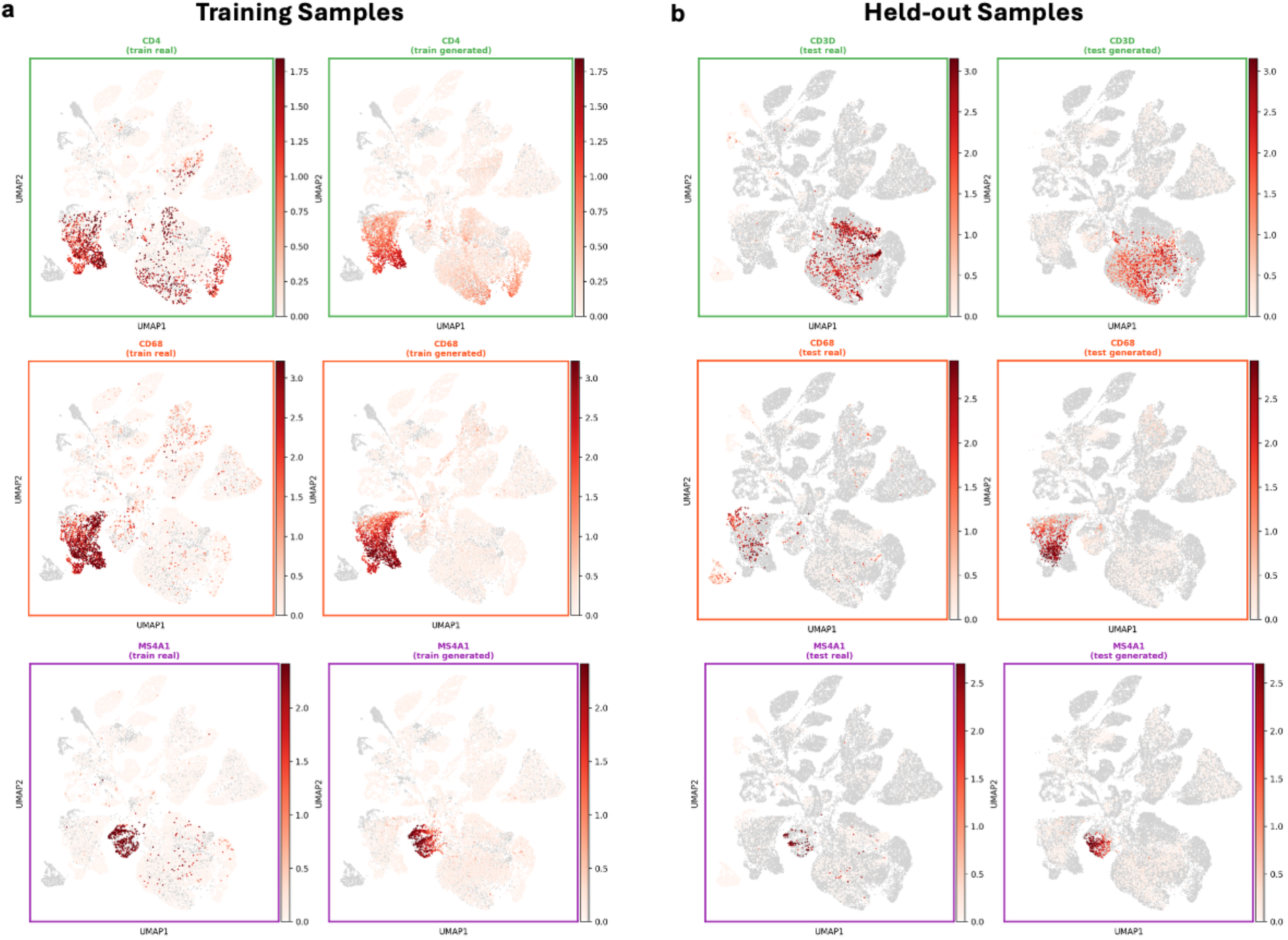
Reconstruction of immune marker gene expression. Expression of *CD3D* (T cells), *CD68* (macrophages), and *MS4A1* (B cells) in the breast cancer dataset is shown for real (left) and generated (right) cells in training (a) and held-out samples (b).

The same behavior was observed in AML, where generated cells reproduced expression of *CD3D*, *CD14*, and *MS4A1* in the expected T-cell, macrophage, and B-cell regions, respectively, across both training and held-out samples (Supplementary Figure 3). Likewise, *ACTB* and *HPRT1* showed comparable expression patterns between real and generated cells in both evaluation settings (Supplementary Figure 4). Taken together, these analyses indicate that marker gene expression and broader baseline expression patterns were reproduced consistently across both cancer types, including in held-out samples.

### Reconstruction of malignant marker gene expression

We then examined whether the same level of agreement extended to malignant marker gene expression, which is expected to exhibit greater inter-sample heterogeneity than immune marker expression. In training samples, generated cells reproduced expression of *ESR1*, associated with hormone receptor-positive cancer cells, and *ERBB2*, associated with HER2-positive cancer cells, in the corresponding cancer cell regions of the embedding (Figure 4a). This indicates that, when similar subtype-associated malignant states were represented during training, the model could generate cancer cell populations with similar marker expression patterns.

**Figure 4:**
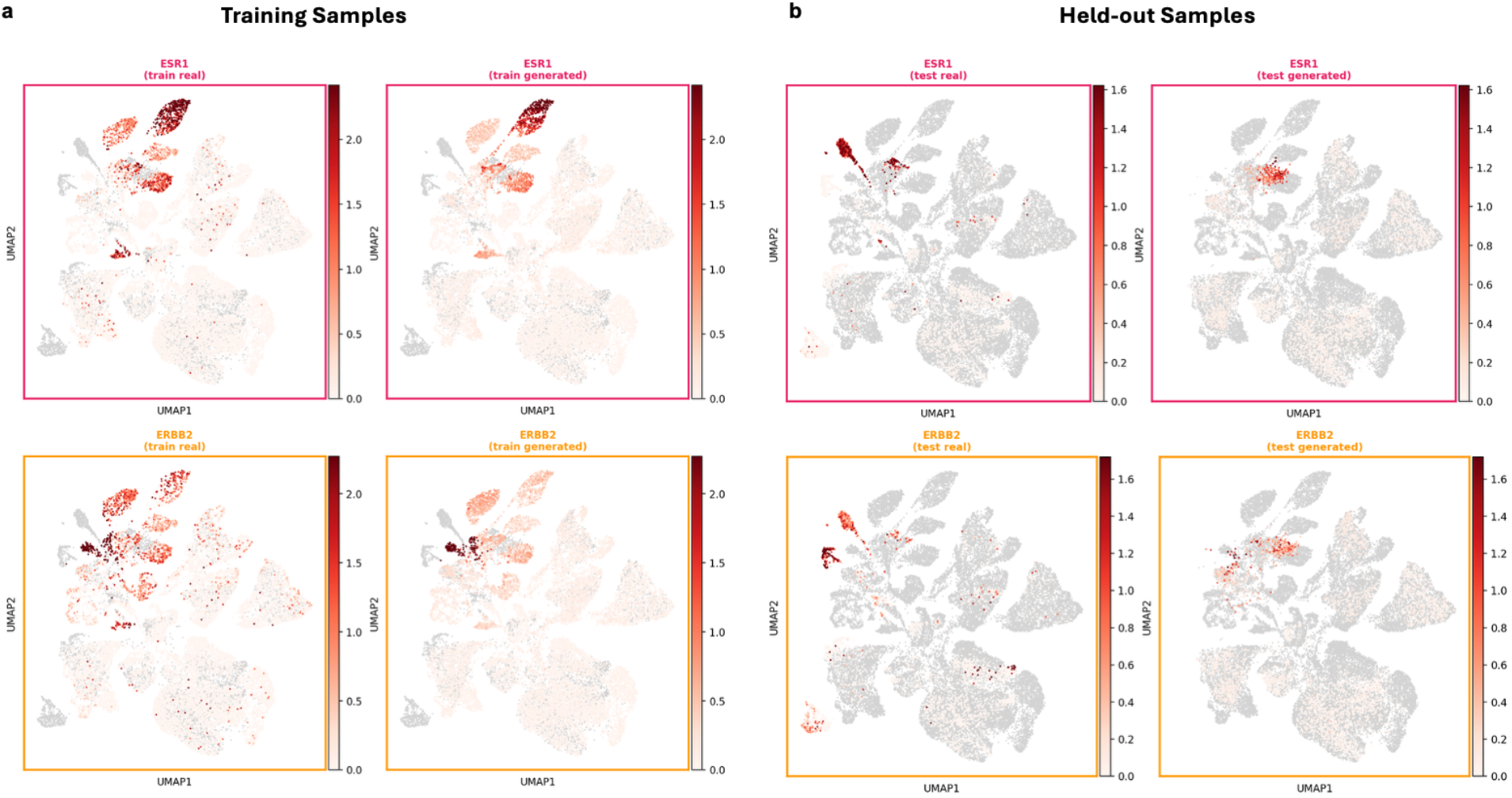
Reconstruction of malignant marker gene expression. Expression of *ESR1* (hormone receptor-positive cancer cells) and *ERBB2* (HER2-positive cancer cells) in the breast cancer dataset is shown for real (left) and generated (right) cells in training (a) and held-out samples (b).

Performance was more inconsistent in held-out breast cancer samples. Some generated malignant populations retained *ESR1* or *ERBB2* expression patterns consistent with the matched real data, particularly in embedding regions that were also well covered by training samples (Figure 4b). However, other held-out malignant cell regions were only partially captured or absent in the generated data. Thus, unlike the more consistent reconstruction of immune marker expression, malignant marker expression generalized less reliably to held-out samples, particularly for sample-specific states that were insufficiently represented in the training cohort.

These findings are consistent with the global UMAP analysis and indicate that bulk2scDiff more reliably reconstructs recurrent biological features shared across samples than sample-specific malignant states under unseen pseudobulk conditions.

### Reconstruction of individual samples

At the individual-sample level, we investigated whether a single pseudobulk profile could generate the cellular landscape of its corresponding sample rather than only preserving pooled dataset-level structure. For the breast cancer cohort, representative training and held-out samples were visualized in their own UMAP spaces, comparing real and generated cells per sample (Figure 5). In training samples, generated cells closely matched the full real-cell distribution, including the major immune and malignant compartments (Figure 5a). This sample-level agreement indicates that bulk2scDiff can generate cellular distributions consistent with a training sample’s pseudobulk profile.

**Figure 5:**
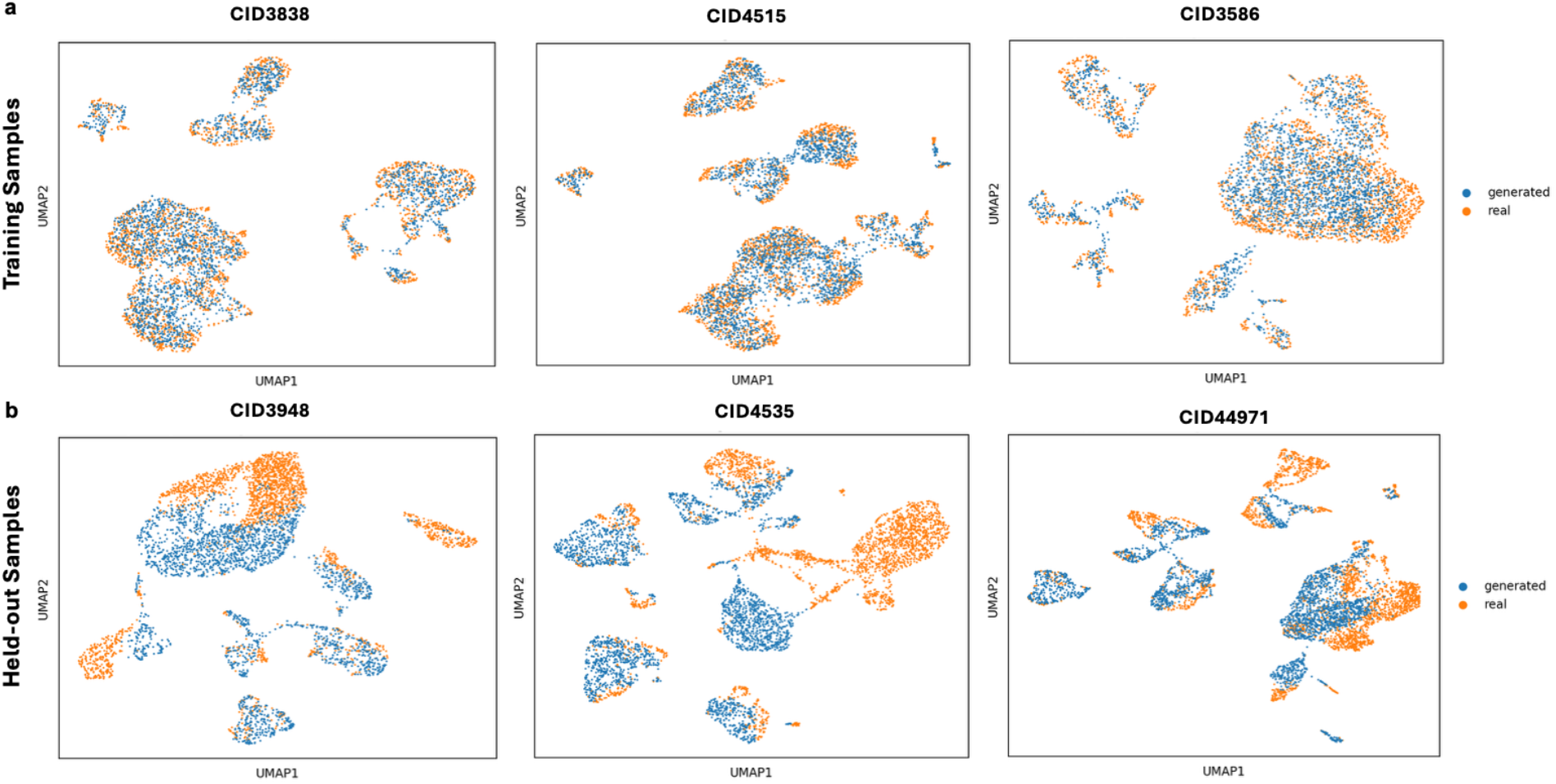
Reconstruction of individual-sample cellular landscapes. Real (orange) and generated (blue) single-cell profiles are shown for three representative training samples (a) and three representative held-out samples (b) from the breast cancer dataset, with each sample visualized in its own UMAP embedding space.

In held-out breast cancer samples, generated populations remained broadly aligned with the real single-cell data but showed more restricted coverage of the cellular manifold (Figure 5b). Most major cell populations were represented, but some sample-specific subpopulations, particularly malignant cell clusters, were reduced or missing. The generated cells therefore captured the dominant sample-level structure while incompletely reconstructing the full diversity of individual held-out tumors. This behavior is consistent with the preceding global and marker-gene analyses: training-sample reconstruction was strong, while held-out generalization was partial and depended on whether the relevant cell states had been sufficiently represented during training.

The same individual-sample analysis in AML supported this conclusion in a second disease context (Supplementary Figure 5). AML training samples were reconstructed closely, whereas held-out AML samples showed good recovery of major cell populations but incomplete coverage of some sample-specific substructures. Thus, similar reconstruction behavior was observed across both cancer types, with the strongest performance for recurrent biological features shared across samples and reduced performance for sample-specific cellular states.

We further assessed that whether generated populations remained quantitatively consistent with the aggregate expression profile used for conditioning. In training samples, per-sample Pearson correlations between each sample’s conditioning pseudobulk and a proxy pseudobulk recomputed from its generated cells were high (breast cancer mean 0.96, range 0.93 to 0.98, n = 21; AML mean 0.95, range 0.91 to 0.97, n = 32). This agreement is expected given that the model is optimized to match the conditioning pseudobulk, and it primarily confirms that the generative pipeline behaves as intended.

The more informative comparison is in held-out samples, where this correlation remained high, with a larger reduction in breast cancer (mean 0.88, range 0.84 to 0.93, n = 5) than in AML (mean 0.93, range 0.90 to 0.96, n = 9) (Supplementary Figure 6). Because held-out samples were not part of the optimization target, this agreement indicates that the generated populations retained substantial correspondence with the aggregate expression profile of samples the model had not been trained to match. However, high agreement at the pseudobulk level alone cannot establish that generation is driven by sample-specific conditioning, because samples within the same cohort share substantial transcriptomic structure. We therefore tested conditioning specificity directly.

### Conditioning specificity by pseudobulk swapping

To directly evaluate whether generated populations were specifically associated with their conditioning pseudobulk rather than reflecting only shared cohort-level structure, we performed a pseudobulk swapping experiment. For each dataset, every real sample was compared against every generated population, so that each real sample had one matched population (generated from its own pseudobulk) and multiple mismatched populations (generated from every other sample’s pseudobulk). We quantified each comparison in latent space using two complementary distances: maximum mean discrepancy (MMD) with a radial basis function (RBF) kernel (22), which is sensitive to local differences in population density, and energy distance (23), based on pairwise Euclidean distances, which is sensitive to differences in overall location and spread. Agreement between the two metrics therefore constitutes convergent rather than redundant evidence. For each real sample and each metric, we ranked all generated populations by their distance from the real population and recorded whether the matched population ranked first.

Top-1 match rates are reported in Supplementary Table 1; Figure 6 and Supplementary Figure 7 display the corresponding distance matrices as a qualitative view of the full pairwise structure. By both metrics, the population generated from a sample’s own pseudobulk was the closest match to that sample’s real cells in nearly every comparison. By MMD, this held for all breast cancer samples, 21 of 21 training and 5 of 5 held-out, and for 40 of 41 AML samples, 31 of 32 training and 9 of 9 held-out. By energy distance, this held for 25 of 26 breast cancer samples, 21 of 21 training and 4 of 5 held-out, and for all 41 AML samples, 32 of 32 training and 9 of 9 held-out (Supplementary Table 1). The only two exceptions, one AML training sample under MMD and one breast cancer held-out sample under energy distance, were near-miss cases rather than clear failures. Because the exceptions occurred independently in a training sample and a held-out sample, they are more consistent with isolated metric-level variation than with a systematic loss of specificity in unseen samples, indicating that the pseudobulk condition carries sample-specific information that is largely preserved for pseudobulk profiles not observed during diffusion training.

**Figure 6:**
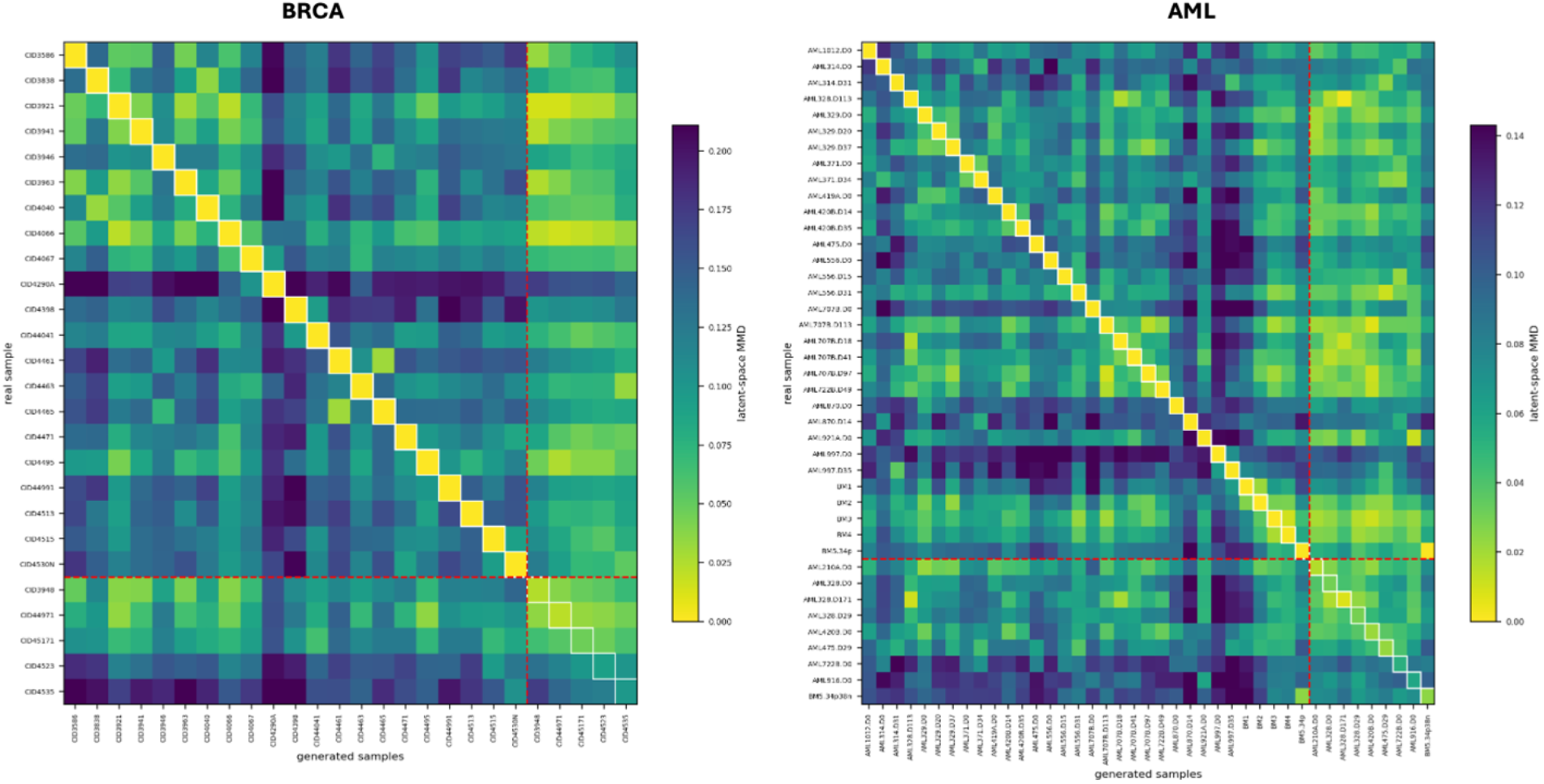
Conditioning specificity by pseudobulk swap. Pairwise latent-space MMD between real samples (rows) and populations generated under each sample’s pseudobulk condition (columns) for BRCA (left) and AML (right). Lower values indicate greater similarity. White outlines denote matched sample condition pairs, and red dashed lines separate training and held-out samples.

## Discussion

This study establishes an initial proof of principle that pseudobulk transcriptomic profiles can condition a diffusion model to generate sample-associated single-cell RNA-seq distributions. Across two cancer datasets, bulk2scDiff closely reconstructed populations from training samples and generated biologically coherent populations for held-out samples, while retaining sample-specific conditioning in nearly all pseudobulk-swap comparisons. These findings support the feasibility of formulating bulk-to-single-cell inference as a conditional generative problem rather than restricting inference to summary quantities such as cell-type proportions or cell-type-averaged expression profiles. At the same time, the reduced reconstruction of sample-specific malignant features in held-out samples defines an important boundary of what can currently be inferred from an aggregate transcriptomic profile.

This boundary is closely related to the underdetermined nature of the problem. Distinct combinations of cell-type abundance and cell-state expression can produce similar or nearly identical aggregate transcriptomic profiles (24,25). Consequently, conditional generation should not be interpreted as recovering a unique ground-truth cellular population from an aggregate measurement. The conditioning profile instead constrains generation, while the single-cell populations that can be generated also depend on the biological variation represented in the training data. An unseen aggregate profile may therefore contain sample-specific information without providing sufficient information to recover cellular states that are poorly represented or absent from the training distribution.

The comparison between training and held-out samples illustrates this limitation. The bulk2scDiff framework closely reconstructed the cellular distributions of training samples but showed weaker and more inconsistent generalization to held-out samples. Recurrent immune features were comparatively well preserved, whereas sample-specific malignant expression patterns showed substantially weaker generalization. This asymmetry is biologically plausible given the substantial inter-sample heterogeneity of malignant populations observed across both cancer contexts (18,19), and indicates that conditional generation generalizes more readily to cellular features that recur across samples than to highly individualized states. The relatively limited number and diversity of independent training samples likely further constrain this generalization, particularly for rare or sample-specific states. Larger and more diverse cohorts will therefore be important for determining how well the framework generalizes across biological variation.

An important limitation is that the conditioning profiles used here were pseudobulk profiles computed from the same scRNA-seq cohorts rather than independently measured bulk RNA-seq. This design provides matched single-cell ground truth and enables controlled evaluation of the relationship between an aggregate profile and its underlying cellular distribution, but pseudobulk and experimentally measured bulk RNA-seq are not equivalent. Differences in tissue dissociation, cell capture, RNA processing, and sequencing protocols can introduce systematic differences in both cellular representation and measured gene expression (26,27). Performance under pseudobulk conditioning should therefore not be assumed to transfer directly to experimentally measured bulk RNA-seq. Validation in cohorts with matched bulk and single-cell measurements will be necessary to determine whether the learned conditional relationship remains robust across this domain shift.

The present evaluation also has methodological limitations. First, bulk2scDiff was not systematically benchmarked against existing approaches that infer cellular composition or expression from aggregate transcriptomic data. Although conventional deconvolution methods do not aim to generate complete single-cell distributions, comparison on shared endpoints would help establish what additional information is provided by conditional generation. Second, the current implementation relies on a single pretrained SCimilarity (20) encoder-decoder to define the latent representation in which diffusion is performed. Alternative single-cell representations could be explored using pretrained foundation models such as scGPT (28) and scFoundation (29), as well as learned latent representations such as scVI (13). Comparing these representations will help determine how strongly performance depends on the choice of latent space.

Overall, bulk2scDiff shows the feasibility of using aggregate transcriptomic information to condition the generation of single-cell populations in a controlled pseudobulk setting. The results support conditional generation as a complementary formulation of bulk-to-single-cell inference, while highlighting that generalization depends on the biological diversity represented during training. Extending this framework to experimentally measured bulk RNA-seq will be an important next step toward evaluating its applicability to clinical bulk transcriptomic cohorts.

## Methods

### Datasets

We evaluated the pseudobulk-conditioned single-cell diffusion framework on two publicly available single-cell RNA-seq datasets representing distinct tumor contexts (Table 1). The datasets comprised a breast cancer cohort from Wu et al. (19) and an acute myeloid leukemia (AML) cohort from van Galen et al. (18). Gene filtering was applied independently within each cohort, so the retained gene space differs between the two datasets.

After preprocessing, the breast cancer model used 25,209 retained genes across 26 samples and 99,526 cells. Because this cohort contains clinically relevant tumor subtypes, the sample-level split was fixed manually to preserve subtype representation, yielding 21 training samples (9 ER-positive, 4 HER2-positive, 8 triple-negative) and 5 held-out samples (2 ER-positive, 1 HER2-positive, 2 triple-negative). The AML model used 19,616 retained genes across 41 samples and 37,808 cells after excluding cell-line samples. The two cell-line samples (MUTZ3 and OCI-AML3) were excluded before splitting. The remaining 41 samples were partitioned at the sample level using an 80/20 split with a fixed random seed of 1234, giving 32 training and 9 held-out samples, with five of the six normal bone marrow samples assigned to training and one to the held-out set.

Sample assignments were fixed before diffusion-model training. Cells from held-out samples were excluded from diffusion training and were used only for post-training conditional generation and downstream evaluation. The autoencoder representation was learned separately, as described below.

### Single-cell preprocessing

Input single-cell count matrices were processed with a common preprocessing procedure before autoencoder encoding, pseudobulk construction, or model evaluation. Raw data were provided in AnnData format. Gene names were made unique, genes detected in fewer than 3 cells were removed, and cells expressing fewer than 10 genes were removed.

For each retained cell, the original library size was recorded from the full raw count matrix before gene filtering; these pre-filter library sizes were then used as the normalization denominators, so that the total sequencing depth of each cell was preserved even after low-detection genes were removed. The count matrix was subset to retained cells and genes, normalized to a target library size of 10,000 using the pre-gene-filter cell library size, and transformed with log1p.

Only minimal count-depth and sparsity filtering was applied. Cells were not filtered according to mitochondrial transcript fraction, doublet score, additional total-UMI thresholds, or marker-based cell-type annotations. The same preprocessing function was used during autoencoder training, diffusion training, sample generation, and evaluation, so that all model components operated in a consistent gene space. Sample-level training and held-out partitions were defined using the sample identifier supplied in the corresponding dataset metadata.

Cell-type labels were taken directly from the metadata provided by the original publications and were not re-annotated, re-clustered, or otherwise modified. These labels were used only for visualization and for interpreting generated populations, and were not used as model input or as a training signal.

### Pseudobulk conditioning vectors

A pseudobulk conditioning vector was constructed independently for each biological sample from the raw count matrix. Raw counts were summed across all cells associated with the same sample identifier within the retained gene space, producing one aggregated expression vector per sample. The corresponding sample-level library size was calculated as the sum of the pre-gene-filter library sizes of all cells assigned to that sample. The aggregated counts were then normalized to a target library size of 10,000 using this sample-level denominator and transformed using log1p.

The resulting vector represents a normalized bulk-like summary of the sample and differs from an average of independently normalized single-cell profiles. Because the gene-filtering mask was shared across the cohort and normalization used pre-filter library-size totals, the procedure was numerically equivalent to aggregating the complete raw count matrix before restricting it to the retained genes.

The pseudobulk vectors were derived from the same scRNA-seq datasets and therefore served as controlled stand-ins for independently measured bulk RNA-seq profiles. This design provided a matched single-cell reference population for each conditioning input but did not model technical differences between independently generated bulk and single-cell measurements. During evaluation, the pseudobulk vector of each target sample was calculated using the same procedure and supplied as the conditioning input.

### Latent representations of single cells and pseudobulk conditioning

The diffusion model operated in a learned low-dimensional latent space rather than directly in gene-expression space. Single-cell expression profiles were encoded using an autoencoder based on the SCimilarity encoder-decoder architecture (20). The encoder contained three fully connected hidden layers of width 1,024, each followed by batch normalization and a PReLU activation, and a final linear projection to 128 dimensions. The resulting embedding was L2-normalized to lie on the unit hypersphere, following the original SCimilarity design. The decoder mirrored the encoder, using three fully connected layers of width 1,024 with batch normalization and PReLU activations, followed by a linear projection to the dataset-specific gene space with no output activation. Dropout was disabled, so latent representations were deterministic. During autoencoder fine-tuning, the reconstruction loss was computed on this unrectified linear output. For all downstream gene-space analyses, decoded profiles were rectified so that negative reconstructed values were set to zero.

The autoencoder was initialized using publicly released SCimilarity weights (model version 1.1). The input layer of the SCimilarity encoder and the output layer of its decoder depend directly on the predefined SCimilarity gene panel and were therefore incompatible with the dataset-specific retained gene sets used here. These two layers were replaced with newly initialized layers matching the number of genes retained in each dataset. The remaining hidden layers do not depend on the input or output gene-space dimensionality and were retained as a pretrained initialization. The complete autoencoder, including both retained and newly initialized layers, was subsequently fine-tuned end to end on each dataset.

Autoencoder fine-tuning minimized the mean squared reconstruction error between the input and reconstructed log-normalized expression profiles. Optimization used AdamW with a learning rate of 5 × 10⁻⁴, weight decay of 0.01, a batch size of 128, and 200,000 gradient steps. The autoencoder was fine-tuned using all cells in each cohort. After fine-tuning, the encoder parameters were frozen and used to project all cells into the 128-dimensional latent representation. The decoder was retained for reconstruction diagnostics and gene-space visualization.

Each biological sample was associated with an independently constructed gene-space pseudobulk vector *p_s_* ∈ ℝ*^G^*, where *G* is the number of retained genes, and *s* denotes the sample identity. During diffusion training, each single-cell latent vector was paired with the pseudobulk vector of its source sample. Thus, all cells from the same sample shared the same pseudobulk condition, whereas cells from different samples, including cells within the same minibatch, retained distinct sample-specific conditioning vectors.

The sample-specific pseudobulk vector was encoded using a separate pseudobulk conditioning network. This network did not share weights with the single-cell autoencoder and was trained jointly with the diffusion model. The input pseudobulk vector was layer-normalized and passed through two fully connected hidden layers of width 512, each followed by a SiLU activation and layer normalization. A final linear layer produced a 128-dimensional conditioning embedding *c_s_*. The conditioning embedding therefore matched the dimensionality of the 128-dimensional single-cell latent representation. However, it represented sample-level transcriptomic context rather than an individual cell state and was not explicitly constrained to occupy the same latent geometry as the single-cell embeddings. Instead, it was supplied to the denoising network as a sample-specific conditioning signal.

### Pseudobulk-conditioned latent diffusion model

The generative model was implemented as a denoising diffusion probabilistic model (DDPM) (16) operating in the 128-dimensional single-cell latent representation. The architecture was adapted from the scDiffusion framework (17), and the overall training and evaluation workflow is summarized in Figure 1.

#### Forward diffusion process

A forward Markov chain progressively corrupts a clean latent vector *z*_0_ over *T* = 1,000 discrete timesteps using a linear variance schedule, with *β*_1_ = 1 × 10^−4^ increasing linearly to *β_T_* = 0.02. A noisy latent at timestep *t* was sampled directly as:

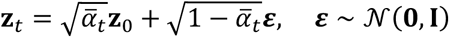

Where

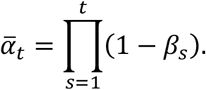

This closed-form expression allowed arbitrary diffusion timesteps to be sampled during training without explicitly simulating all preceding forward transitions.

#### Denoising network and conditioning

The denoising network *ε_θ_*(*z_t_*, *t*, *c_s_*) is a fully connected latent U-Net with residual blocks, hidden widths of 512, 512, 256, and 128 units, and skip connections from the encoder path to the decoder path. The network was conditioned jointly on the diffusion timestep *t* and the sample-specific pseudobulk conditioning embedding *c_s_*.

The timestep was represented using a sinusoidal embedding followed by a two-layer multilayer perceptron with a SiLU activation. Pseudobulk conditioning entered the denoising network through two mechanisms. First, the pseudobulk embedding *c_s_* was projected into the timestep-embedding space and added to the timestep representation. Second, *c_s_* was used to generate feature-wise scale and shift parameters for FiLM (21) within each residual block. The scale and shift parameters were produced by a linear projection of *c_s_* within each residual block.

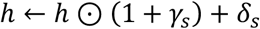

The sample-specific conditioning embedding therefore influenced both the global timestep-conditioning pathway and the transformations performed throughout the residual network. The final network output was projected to 128 dimensions and interpreted as the predicted Gaussian noise contained in the noisy single-cell latent vector.

Conceptually, the model estimates a conditional distribution of single-cell latent states given a sample-level pseudobulk profile. Repeated reverse-diffusion sampling under the same sample-specific pseudobulk condition can therefore generate a population of synthetic single-cell latent profiles rather than a single deterministic reconstruction.

### Diffusion model training

The diffusion model was trained using latent representations of cells from the training samples only. At each step, a batch of clean single-cell latent vectors *z*_0_ was sampled. Each latent vector was paired with the conditioning embedding c*_s_*_(*i*)_ derived from the pseudobulk profile of its source sample *s*(*i*). A timestep *t* ∼ Uniform{1, …, 1000} was sampled per example, and Gaussian noise *ε* was used to construct *z_t_* via the forward diffusion process. The model was trained using the standard noise-prediction objective:

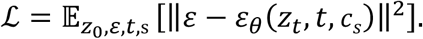

Optimization used AdamW with a learning rate of 1 × 10⁻⁴, weight decay of 1 × 10⁻⁴, and a batch size of 128, with the learning rate annealed linearly to zero over 1,000,000 gradient steps.

An exponential moving average of the model parameters was maintained during training with a decay of 0.9999, but all reported results were generated using the final unaveraged parameter checkpoint. Checkpoints were saved every 200,000 steps. The final checkpoint corresponded to the predefined training budget and was not selected according to held-out-sample performance. This fixed-budget criterion was used so that no held-out-sample information entered model selection, at the cost of not optimizing the checkpoint with respect to generation quality. One training run was performed per dataset; replicate runs with different random initializations were not conducted.

Optimization stability was assessed from the training loss and associated training diagnostics. Training-sample MMD was additionally calculated at predefined checkpoints to examine how closely generated and observed training-sample latent distributions converged during model fitting. These training-only trajectories are shown in Supplementary Figure 8 and were treated as convergence diagnostics rather than as an estimate of held-out generalization.

### Conditional generation and decoding

For conditional generation, the gene-space pseudobulk vector p*_s_* of a target sample s was calculated using the same aggregation and normalization procedure as during training and transformed by the trained pseudobulk encoder into the corresponding 128-dimensional conditioning embedding *c_s_*. Generation began from isotropic Gaussian noise *z_T_* ∼ N(0, *I*) and proceeded through the learned reverse-diffusion transitions over all 1,000 timesteps. The reverse transitions used a fixed variance schedule rather than a learned variance, with the variance at timestep *t* set to *β_t_* for *t* ≥ 1 and to the corresponding posterior variance at *t* = 1. Each reverse-diffusion trajectory produced one synthetic single-cell latent vector. Multiple trajectories under a shared condition were used to generate a sample-specific synthetic cell population.

For paired comparisons, the number of generated cells was set to the number of available real cells in the corresponding sample, thereby controlling for differences in population size during evaluation. Generated latent vectors were transformed to gene space using the trained autoencoder decoder and rectified as described above. Decoded profiles were used for visualization, pseudobulk-agreement analyses, and marker-gene diagnostics, whereas diffusion training and the primary distributional comparisons were performed in latent space.

### Training and evaluation procedure

After completion of model training, the fixed diffusion model was applied separately to training and held-out samples without further parameter updates. For each evaluated sample, its pseudobulk vector was calculated from raw counts using the training-time preprocessing procedure, and reverse diffusion was performed under the resulting fixed condition. Training-sample generation was used to characterize model fit, whereas generation for held-out samples was used to assess conditional transfer to sample conditions not observed during diffusion-model training. Because the autoencoder was fine-tuned on the complete cohort, the held-out analysis evaluates generalization of the conditional diffusion component rather than fully end-to-end generalization of all learned model components.

### Evaluation metrics and diagnostics

The primary quantitative comparison between real and generated single-cell populations was performed in the shared autoencoder latent representation using radial-basis-function maximum mean discrepancy (RBF-MMD) (22). For each sample, encoded real cells were compared with synthetic latent cells generated under that sample’s pseudobulk condition. Before computing MMD, the real and generated latent populations of each sample were independently subsampled to at most 1,200 cells using a fixed random seed of 1234, so that comparisons were made at comparable population sizes. The kernel bandwidth σ was determined by the median heuristic applied to the pooled real and generated latent vectors, taken as the square root of the median of the non-zero pairwise squared Euclidean distances. Lower MMD values indicate greater similarity between the two latent distributions.

Training-sample MMD trajectories calculated during optimization were used only as training diagnostics and are reported in Supplementary Figure 8. Held-out samples were not used for checkpoint selection or other model updates.

To assess whether generated populations were specifically associated with their corresponding pseudobulk conditions and not with a common cohort-level distribution, we performed a conditioning-specificity pseudobulk-swap analysis. For each dataset, the real population of every sample was compared with the synthetic populations generated under every available sample condition, producing a sample-by-condition distance matrix. Each real sample therefore had one matched comparison, corresponding to generation under its own pseudobulk condition, and *N* − 1 mismatched comparisons, corresponding to generation under other samples’ conditions. Real and generated cells were encoded into the shared autoencoder latent space and subsampled using the same 1,200-cell cap and fixed random seed described above.

Distances were quantified using RBF-MMD and, as an alternative distributional metric, latent-space energy distance (23), as previously applied to single-cell population comparisons in scPerturb (30). The energy distance was computed directly from pairwise Euclidean distances between latent vectors rather than through an external package. For each real sample, the *N* distances in the corresponding row of the matrix were ranked in ascending order and the rank of the matched comparison was recorded. A sample was scored as a top-1 match when the matched comparison gave the smallest distance among all candidate conditions, and the reported counts represent the number of samples meeting this criterion. Summary statistics over mismatched comparisons were calculated after excluding the diagonal of the matrix. Because the unbiased squared-MMD estimator is not constrained to be non-negative, matched MMD values for closely reproduced populations were occasionally slightly negative; these values were used as returned by the estimator and were not truncated.

Per-sample distances, ranks and top-1 assignments were exported to a summary table, and the resulting match rates are reported in Supplementary Table 1. The distance matrices themselves are shown as heatmaps with training and held-out samples annotated separately (Figure 6 for MMD and Supplementary Figure 7 for energy distance); these panels illustrate the matched-versus-mismatched contrast qualitatively and are not the source of the reported counts. The heatmaps visualize the complete pairwise distance structure, whereas the reported top-1 counts were derived computationally from the row-wise rankings of the underlying numerical matrices.

Additional qualitative and biological diagnostics were used to characterize model behavior. Joint UMAP embeddings were generated with Scanpy (31). For dataset-level analyses, a single joint embedding was computed per dataset and reused across the corresponding global and marker-gene panels so that coordinates were directly comparable. Individual-sample analyses were visualized using sample-specific UMAP embeddings containing the corresponding real and generated cells. Sample-level agreement was further examined by comparing each input pseudobulk vector with a proxy aggregate calculated from the corresponding decoded synthetic population. Marker-gene analyses compared the spatial distribution and expression intensity of selected immune and tumor-associated genes between real and generated populations in the shared embedding. These analyses were interpreted as complementary biological diagnostics rather than as independent evidence of exact single-cell reconstruction.

### Software and implementation

All analyses were performed in Python 3.9 using PyTorch 1.13.0, numpy 1.23.4, pandas 1.5.1, scikit-learn 1.2.2, anndata 0.8.0, Scanpy 1.9.1 (30), and umap-learn 0.5.7. Random seeds were fixed for autoencoder fine-tuning, diffusion-model training, conditional generation, and all downstream subsampling; the specific values used for each stage and dataset are recorded in the released code repository. Model training was performed on a single NVIDIA GeForce RTX 4080 SUPER GPU, with approximately six hours required for 1,000,000 diffusion training steps.

## Data and code availability

The single-cell RNA-seq datasets analyzed in this study are available from the Gene Expression Omnibus under accession GSE116256 for the AML cohort (18) and accession GSE176078 for the breast cancer cohort (19). The pretrained SCimilarity model weights used to initialize the autoencoder (model version 1.1) are available from Zenodo (https://doi.org/10.5281/zenodo.10685499). The code used for preprocessing, model training, sample generation, and evaluation will be made freely available at: https://github.com/raue-lab/bulk2scDiff.git

## Author contributions

Conceptualization: J.X. and A.R. Methodology: J.X. and A.R. Analyses: J.X. and A.R. Visualization: J.X. and A.R. Supervision: A.R. Writing: J.X. and A.R.

## Supplementary Information

Additional file 1: Supplementary Material (Table S1, Figure S1-S8).

## Use of artificial intelligence (AI)

The authors used ChatGPT version 5.5 (OpenAI) and Claude Sonnet 5 (Anthropic) for writing code and editing manuscript text. The authors reviewed all AI-generated content and take full responsibility for it.

## References

1. Puram SV, Tirosh I, Parikh AS, Patel AP, Yizhak K, Gillespie S, et al. Single-Cell Transcriptomic Analysis of Primary and Metastatic Tumor Ecosystems in Head and Neck Cancer. Cell. 2017 Dec 14;171(7):1611–1624.e24. doi:10.1016/j.cell.2017.10.044 PubMed PMID: 29198524.

2. Stuart T, Satija R. Integrative single-cell analysis. Nat Rev Genet. 2019 May;20(5):257–72. doi:10.1038/s41576-019-0093-7

3. Azizi E, Carr AJ, Plitas G, Cornish AE, Konopacki C, Prabhakaran S, et al. Single-cell Map of Diverse Immune Phenotypes in the Breast Tumor Microenvironment. Cell. 2018 Aug 23;174(5):1293–1308.e36. doi:10.1016/j.cell.2018.05.060 PubMed PMID: 29961579; PubMed Central PMCID: PMC6348010.

4. Tirosh I, Izar B, Prakadan SM, Wadsworth MH, Treacy D, Trombetta JJ, et al. Dissecting the multicellular ecosystem of metastatic melanoma by single-cell RNA-seq. Science. 2016 Apr 8;352(6282):189–96. doi:10.1126/science.aad0501

5. Haque A, Engel J, Teichmann SA, Lönnberg T. A practical guide to single-cell RNA-sequencing for biomedical research and clinical applications. Genome Med. 2017 Aug 18;9(1):75. doi:10.1186/s13073-017-0467-4

6. Weinstein JN, Collisson EA, Mills GB, Shaw KRM, Ozenberger BA, Ellrott K, et al. The Cancer Genome Atlas Pan-Cancer analysis project. Nat Genet. 2013 Oct;45(10):1113–20. doi:10.1038/ng.2764

7. Lonsdale J, Thomas J, Salvatore M, Phillips R, Lo E, Shad S, et al. The Genotype-Tissue Expression (GTEx) project. Nat Genet. 2013 Jun;45(6):580–5. doi:10.1038/ng.2653

8. Avila Cobos F, Vandesompele J, Mestdagh P, De Preter K. Computational deconvolution of transcriptomics data from mixed cell populations. Bioinformatics. 2018 Jun 1;34(11):1969–79. doi:10.1093/bioinformatics/bty019

9. Newman AM, Steen CB, Liu CL, Gentles AJ, Chaudhuri AA, Scherer F, et al. Determining cell type abundance and expression from bulk tissues with digital cytometry. Nat Biotechnol. 2019 Jul;37(7):773–82. doi:10.1038/s41587-019-0114-2

10. Wang X, Park J, Susztak K, Zhang NR, Li M. Bulk tissue cell type deconvolution with multi-subject single-cell expression reference. Nat Commun. 2019 Jan 22;10(1):380. doi:10.1038/s41467-018-08023-x

11. Chen Y, Wang Y, Chen Y, Cheng Y, Wei Y, Li Y, et al. Deep autoencoder for interpretable tissue-adaptive deconvolution and cell-type-specific gene analysis. Nat Commun. 2022 Nov 8;13(1):6735. doi:10.1038/s41467-022-34550-9

12. Zhao T, Liu R, Sun Y, Wang B, Zhang L, Chen Q, et al. DECODE: deep learning-based common deconvolution framework for various omics data. Nat Methods. 2026 Mar;23(3):596–608. doi:10.1038/s41592-026-03007-y

13. Lopez R, Regier J, Cole MB, Jordan MI, Yosef N. Deep generative modeling for single-cell transcriptomics. Nat Methods. 2018 Dec;15(12):1053–8. doi:10.1038/s41592-018-0229-2

14. Marouf M, Machart P, Bansal V, Kilian C, Magruder DS, Krebs CF, et al. Realistic in silico generation and augmentation of single-cell RNA-seq data using generative adversarial networks. Nat Commun. 2020 Jan 9;11(1):166. doi:10.1038/s41467-019-14018-z

15. Klein D, Fleck JS, Bobrovskiy D, Zimmermann L, Becker S, Palma A, et al. CellFlow enables generative single-cell phenotype modeling with flow matching [Internet]. bioRxiv; 2025 [cited 2026 Jul 4]. p. 2025.04.11.648220. Available from: https://www.biorxiv.org/content/10.1101/2025.04.11.648220v1 doi:10.1101/2025.04.11.648220

16. Ho J, Jain A, Abbeel P. Denoising Diffusion Probabilistic Models [Internet]. arXiv; 2020 [cited 2025 Dec 8]. Available from: http://arxiv.org/abs/2006.11239 doi:10.48550/arXiv.2006.11239

17. Luo E, Hao M, Wei L, Zhang X. scDiffusion: conditional generation of high-quality single-cell data using diffusion model. Bioinformatics. 2024 Sep 1;40(9):btae518. doi:10.1093/bioinformatics/btae518

18. van Galen P, Hovestadt V, Wadsworth II MH, Hughes TK, Griffin GK, Battaglia S, et al. Single-Cell RNA-Seq Reveals AML Hierarchies Relevant to Disease Progression and Immunity. Cell. 2019 Mar 7;176(6):1265–1281.e24. doi:10.1016/j.cell.2019.01.031

19. Wu SZ, Al-Eryani G, Roden DL, Junankar S, Harvey K, Andersson A, et al. A single-cell and spatially resolved atlas of human breast cancers. Nat Genet. 2021 Sep;53(9):1334–47. doi:10.1038/s41588-021-00911-1 PubMed PMID: 34493872; PubMed Central PMCID: PMC9044823.

20. Heimberg G, Kuo T, DePianto DJ, Salem O, Heigl T, Diamant N, et al. A cell atlas foundation model for scalable search of similar human cells. Nature. 2025 Feb;638(8052):1085–94. doi:10.1038/s41586-024-08411-y

21. Perez E, Strub F, Vries H de, Dumoulin V, Courville A. FiLM: Visual Reasoning with a General Conditioning Layer [Internet]. arXiv; 2017 [cited 2026 Jun 9]. Available from: http://arxiv.org/abs/1709.07871doi:10.48550/arXiv.1709.07871

22. Gretton A, Borgwardt KM, Rasch MJ, Schölkopf B, Smola A. A Kernel Two-Sample Test. J Mach Learn Res. 2012;13(25):723–73.

23. Rizzo ML, Székely GJ. Energy distance. WIREs Comput Stat. 2016;8(1):27–38. doi:10.1002/wics.1375

24. Newman AM, Steen CB, Liu CL, Gentles AJ, Et. A. Determining cell type abundance and expression from bulk tissues with digital cytometry. Nat Biotechnol. 2019. doi:10.1038/s41587-019-0114-2

25. Avila Cobos F, Alquicira-Hernandez J, Powell JE, Mestdagh P, De Preter K. Benchmarking of cell type deconvolution pipelines for transcriptomics data. Nat Commun. 2020 Nov 6;11(1):5650. doi:10.1038/s41467-020-19015-1

26. Hippen AA, Omran DK, Weber LM, Jung E, Drapkin R, Doherty JA, et al. Performance of computational algorithms to deconvolve heterogeneous bulk ovarian tumor tissue depends on experimental factors. Genome Biol. 2023 Oct 20;24:239. doi:10.1186/s13059-023-03077-7 PubMed PMID: 37864274; PubMed Central PMCID: PMC10588129.

27. Cobos FA, Panah MJN, Epps J, Long X, Man TK, Chiu HS, et al. Effective methods for bulk RNA-seq deconvolution using scnRNA-seq transcriptomes. Genome Biol. 2023 Aug 1;24(1):177. doi:10.1186/s13059-023-03016-6

28. Cui H, Wang C, Maan H, Pang K, Luo F, Duan N, et al. scGPT: toward building a foundation model for single-cell multi-omics using generative AI. Nat Methods. 2024 Aug;21(8):1470–80. doi:10.1038/s41592-024-02201-0

29. Hao M, Gong J, Zeng X, Liu C, Guo Y, Cheng X, et al. Large-scale foundation model on single-cell transcriptomics. Nat Methods. 2024 Aug;21(8):1481–91. doi:10.1038/s41592-024-02305-7

30. Peidli S, Green TD, Shen C, Gross T, Min J, Garda S, et al. scPerturb: harmonized single-cell perturbation data. Nat Methods. 2024 Mar;21(3):531–40. doi:10.1038/s41592-023-02144-y

31. Wolf FA, Angerer P, Theis FJ. SCANPY: large-scale single-cell gene expression data analysis. Genome Biol. 2018 Feb 6;19:15. doi:10.1186/s13059-017-1382-0 PubMed PMID: 29409532; PubMed Central PMCID: PMC5802054.

