## Additional file 1: Supplementary Material (Table S1, Figure S1-S8). for "bulk2scDiff: A Pseudobulk-Conditioned Diffusion Model for Bulk-to-Single-Cell RNA-Seq Generation"

**Supplementary Table 1. Pseudobulk conditioning specificity.** For each real sample, distances to populations generated under all pseudobulk conditions were ranked, with a top-1 match indicating that the sample's own pseudobulk condition yielded the smallest distance. Top-1 match rates are reported using MMD and energy distance for all, training, and held-out samples.

| Dataset | Metric | N | Top-1 (all) | Train | Held-out |
|---|---|---|---|---|---|
| BRCA | MMD | 26 | 26/26 (100%) | 21/21 | 5/5 |
| BRCA | E-distance | 26 | 25/26 (96.2%) | 21/21 | 4/5 |
| AML | MMD | 41 | 40/41 (97.6%) | 31/32 | 9/9 |
| AML | E-distance | 41 | 41/41 (100%) | 32/32 | 9/9 |

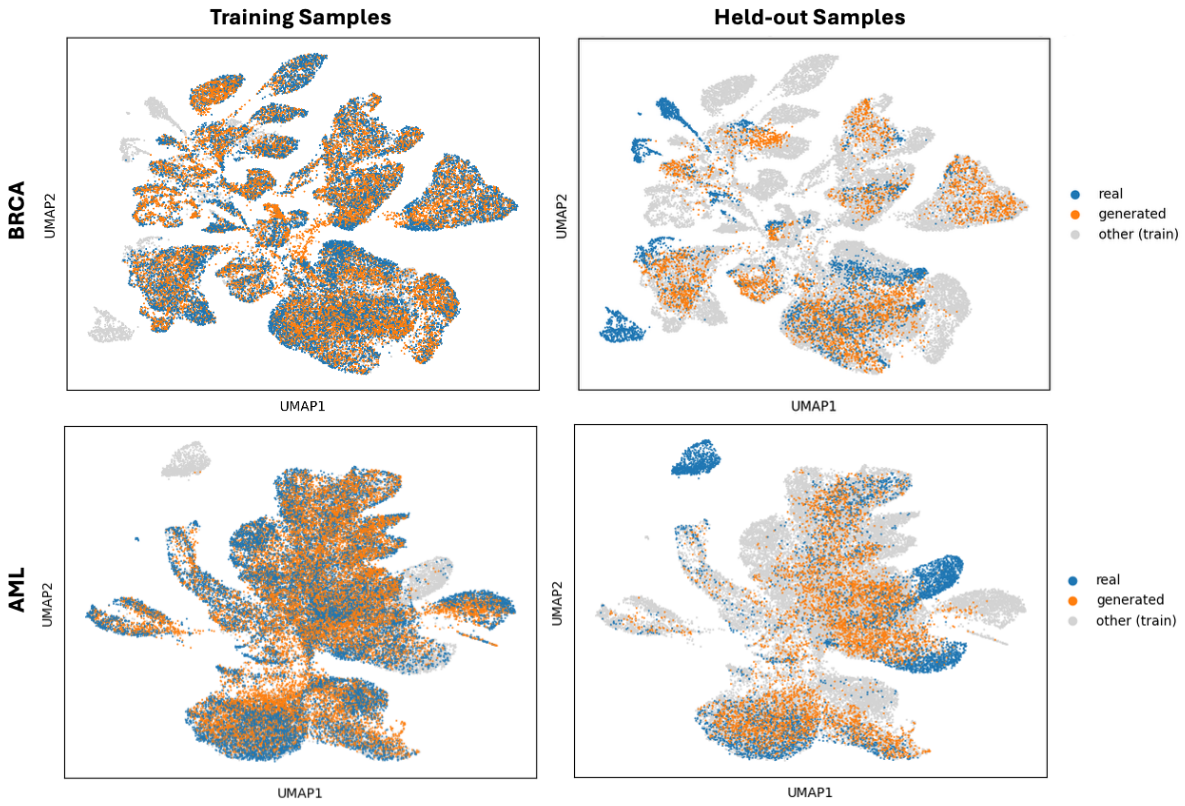


**Supplementary Figure 1: Global performance in training and held-out test samples.** Real and generated cells are shown separately for training (left) and held-out samples (right) in the BRCA (top) and AML (bottom) datasets, using the same UMAP embedding as Figure 2. Gray cells indicate training cells from the remaining samples.

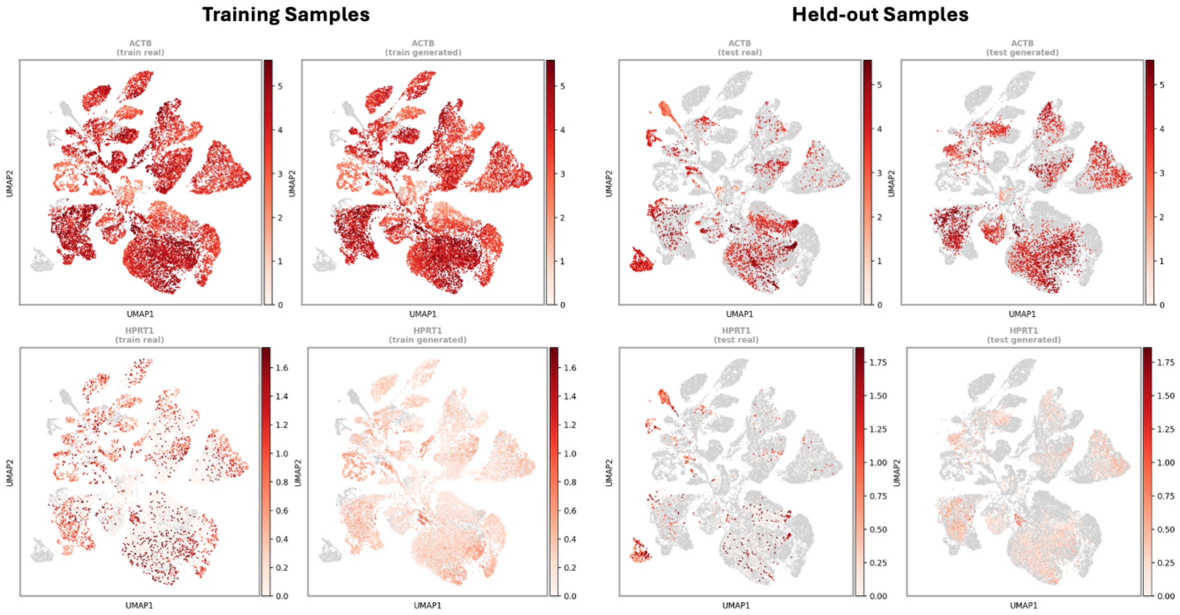


**Supplementary Figure 2: Housekeeping gene expression in BRCA.** Expression of *ACTB* and *HPRT1* is shown for real and generated cells from training and held-out samples using the same UMAP embedding as Figure 2. *ACTB* represents a broadly expressed gene, whereas *HPRT1* represents a lower-expression housekeeping gene.

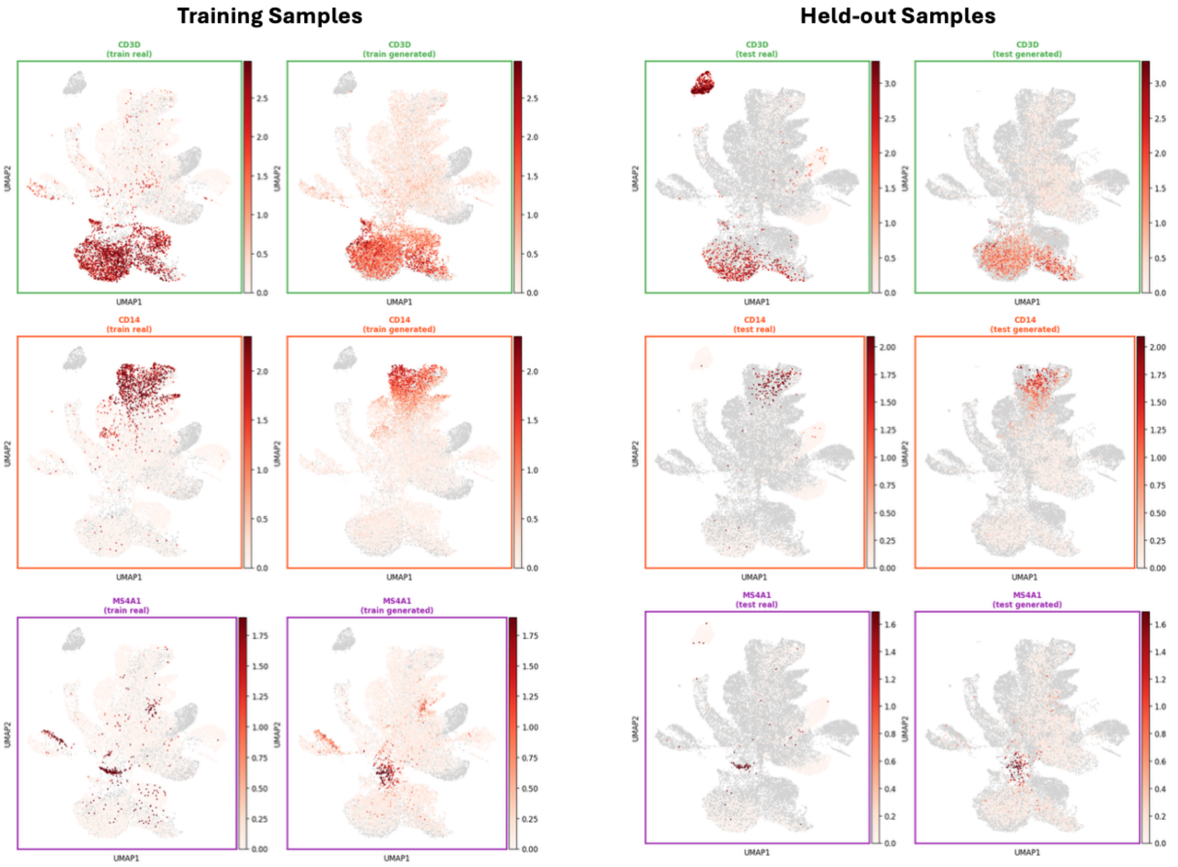


**Supplementary Figure 3: Immune marker gene expression in AML.** Expression of *CD3D*, *CD14*, and *MS4A1*, marking T cells, monocyte/macrophage populations, and B cells, respectively, is shown for real and generated cells from training and held-out samples using the same UMAP embedding as Figure 2.

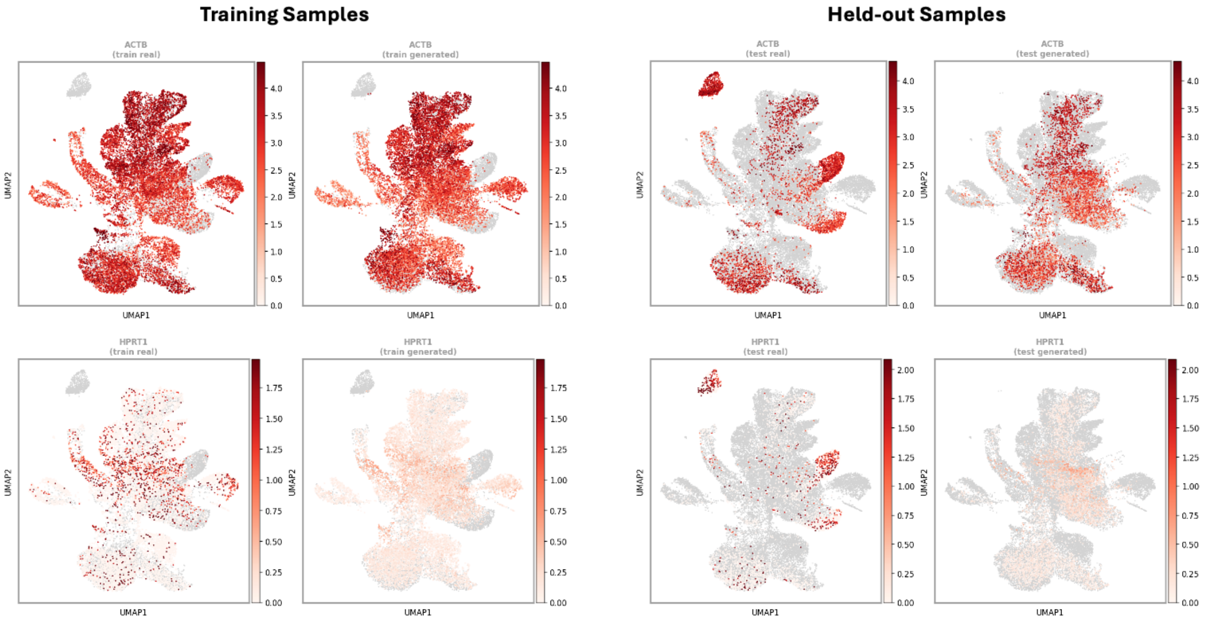


**Supplementary Figure 4: Housekeeping gene expression in AML.** Expression of *ACTB* and *HPRT1* is shown for real and generated cells from training and held-out samples using the same UMAP embedding as Figure 2. *ACTB* represents a broadly expressed gene, whereas *HPRT1* represents a lower-expression housekeeping gene.

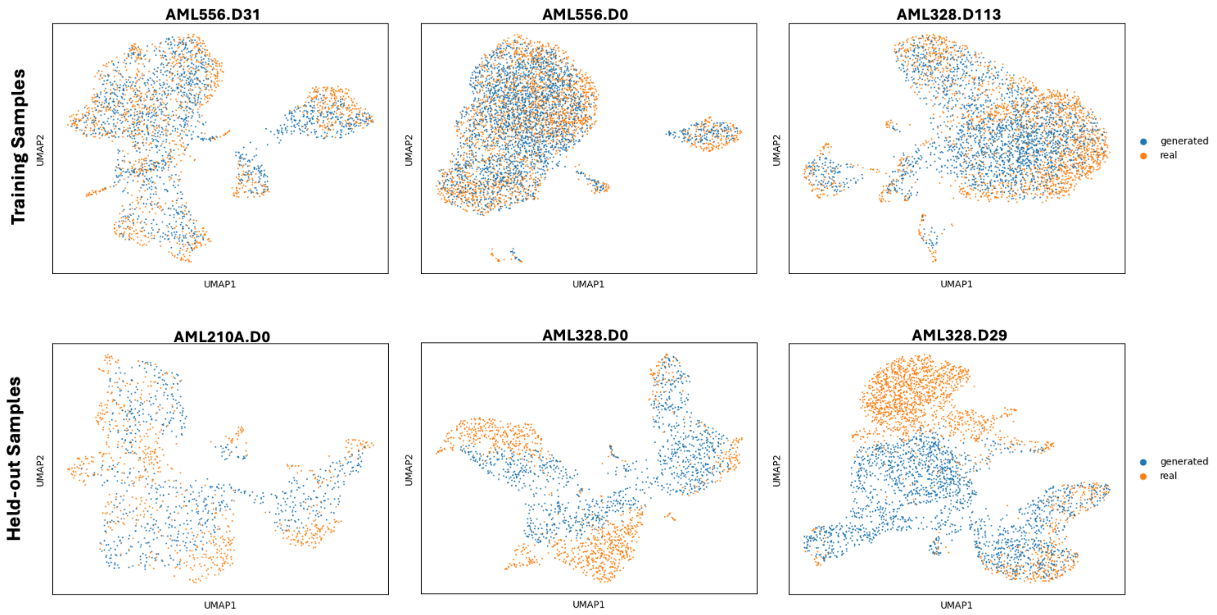


**Supplementary Figure 5: Individual-sample cellular distributions in AML.** Three representative training samples (top) and three representative held-out samples (bottom) are shown, with real and generated cells compared in sample-specific UMAP embeddings.

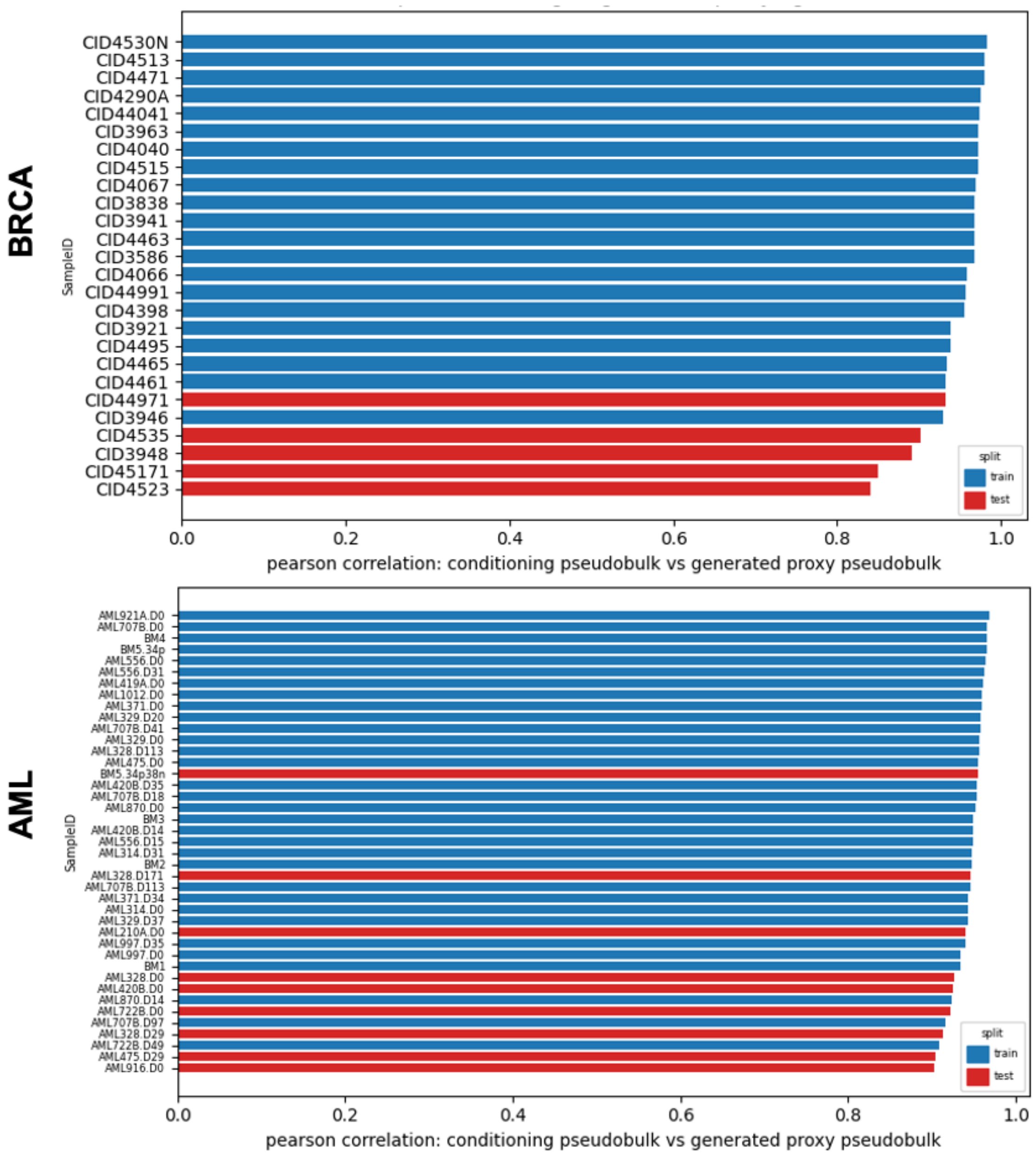


**Supplementary Figure 6: Pseudobulk correlation in training and held-out samples.** Per-sample Pearson correlations compare the conditioning pseudobulk profiles with pseudobulk profiles aggregated from generated cells for BRCA (top) and AML (bottom). Training and held-out samples are shown in blue and red, respectively.

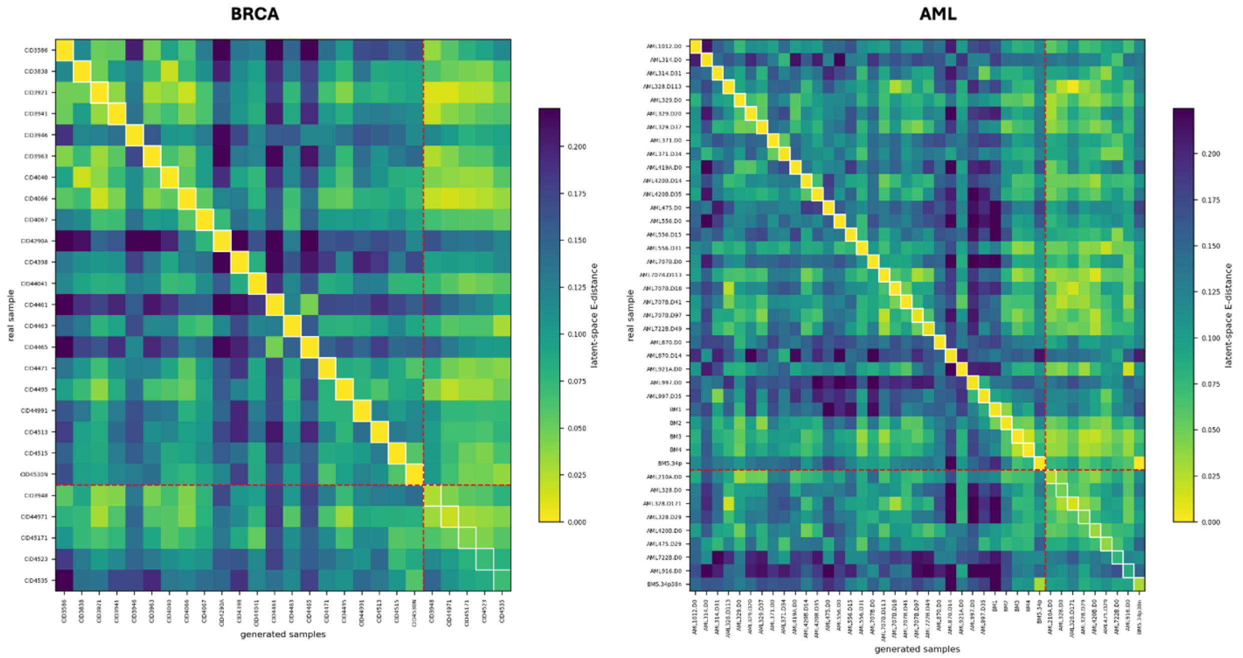


**Supplementary Figure 7: Conditioning specificity by pseudobulk swap (energy distance).** Pairwise latent-space energy distances between each real sample (rows) and populations generated under each sample-level pseudobulk condition (columns) for BRCA (left) and AML (right). Lower values indicate greater similarity. White outlines denote matched sample condition pairs, and red dashed lines separate training and held-out samples.

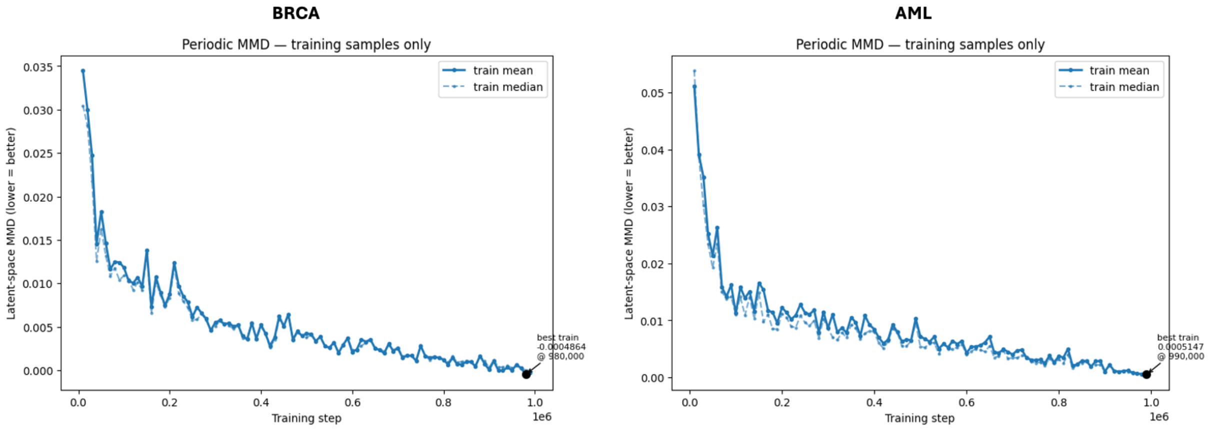


**Supplementary Figure 2: Training dynamics.** Latent-space MMD between generated and corresponding real-cell distributions was evaluated periodically during training for BRCA (left) and AML (right). Solid and dashed lines show the mean and median MMD, respectively. Main-text results use the checkpoint at 1,000,000 training steps.
